# Legumain is a paracrine regulator of osteoblast differentiation and mediates the inhibitory effect of TGF-β1 on osteoblast maturation

**DOI:** 10.1101/2024.04.12.589173

**Authors:** Karl Martin Forbord, Ngoc Nguyen Lunde, Tatjana Bosnjak, Harald Thidemann Johansen, Rigmor Solberg, Abbas Jafari

**Affiliations:** Section for Pharmacology and Pharmaceutical Biosciences, Department of Pharmacy, University of Oslo, 0316 Oslo, Norway; Department of Cellular and Molecular Medicine, University of Copenhagen, 2200 Copenhagen, Denmark

## Abstract

Transforming growth factor-beta 1 (TGF-β1) is a critical regulator of skeletal homeostasis and has diverse promoting effects on osteoblastogenesis. However, the mechanisms behind the intriguing inhibitory effect of TGF-β1 on osteoblast maturation are not fully understood. Here, we demonstrate a novel mechanism by which TGF-β1 modulates osteoblast maturation through lysosomal protease legumain. We observed that presence of TGF-β1 in osteogenic cultures of human bone marrow derived mesenchymal stromal (stem) cells enhanced legumain activity and secretion, in-spite of decreased legumain mRNA expression, suggesting post-transcriptional regulation. We further showed that osteogenic cells internalize and activate prolegumain, associated with inhibited osteoblast maturation, revealing legumain as a paracrine regulator of osteoblast maturation. Interestingly, TGF-β1 treatment exacerbated legumain internalization and activity, and showed an additive effect on legumain-induced inhibition of osteoblast maturation. Importantly, legumain inhibition abolished the inhibitory effect of TGF-β1 on osteoblast maturation. Our findings reveal that TGF-β1 inhibits osteoblast maturation through stimulating secretion and activity of endogenous legumain, as well as increasing the internalization and activation of extracellular legumain. Therefore, our study provides a deeper understanding of the complex regulation of osteoblastogenesis and unveils a novel TGF-β1-legumain axis in regulation of osteoblast maturation and offer novel insights for possible therapeutic interventions related to bone diseases associated with aberrant TGF-β1 signaling.

## 2. Introduction

Transforming growth factor-beta 1 (TGF-β1) is one of the most abundant cytokines in the bone matrix (Hering et al., 2001, Crane and Cao, 2014). This multifunctional cytokine plays a key role in regulation of skeletal homeostasis by orchestrating various cellular processes involved in bone development, remodeling, and repair, which are elucidated by compelling evidence from human and mouse studies (Wu et al., 2024). Domain-specific heterozygous mutations in TGF-β1 in humans are linked to Camurati-Engelmann disease; an autosomal dominant disease associated with increased bone remodeling and osteosclerotic lesions in the long bones and skull (Kinoshita et al., 2000). TGF-β1 deficiency in mice leads to early death due to organ failure associated with autoimmune disease (Kulkarni et al., 1993). Analysis of the tibiae from TGF-β1 deficient mice exhibited decreased longitudinal growth and bone mineral content (Geiser et al., 1998). In addition, histological analysis of bones from TGF-β1 deficient mice indicated depletion of pre-osteoblasts at the trabecular bone surfaces and the presence of osteoprogenitors in the middle of the bone marrow (Atti et al., 2002). At the cellular level, TGF-β1 has been shown to enhance osteoblast proliferation (Kassem et al., 2000) and migration of osteogenic cells to the bone forming surfaces (Tang et al., 2009), while exerting an inhibitory effect on osteoblast apoptosis (Jilka et al., 1998). Furthermore, TGF-β1 has also been shown to enhance commitment of bone marrow stromal cells to the osteogenic lineage (i.e. early-stage osteoblast differentiation) (Wu et al., 2024). In contrast, TGF-β1 also exerts an inhibitory effect on osteoblast maturation (i.e. late-stage osteoblast differentiation) (Maeda et al., 2004). Although the mechanisms behind pro-osteogenic actions of TGF-β1 are well-established and have been shown to be mediated through e.g. stimulating the expression of master regulators of osteogenesis, such as Runx2 and osterix/Sp7 (Seo and Serra, 2009), the mechanism responsible for the intriguing inhibitory effect of TGF-β1 on osteoblast maturation are not well understood.

Legumain (also known as asparaginyl endopeptidase) is a lysosomal cysteine protease involved in protein activation, processing, and degradation, thereby playing roles in diverse biological processes (Solberg et al., 2022). Legumain is also secreted and is measurable in body fluids such as serum/plasma and is thus postulated to have autocrine/paracrine functions (reviewed in (Lunde et al., 2019)). In addition, we have previously shown that high levels of legumain is present in the bone microenvironment (Jafari et al., 2017). Furthermore, we showed that legumain inhibits osteoblast maturation and bone formation by bone marrow mesenchymal stem (stromal) cells (BMSC) through degradation of the extracellular matrix protein fibronectin (Jafari et al., 2017).

Herein, we provide evidence for a novel cross-regulatory function of TGF-β1 and legumain, in which TGF-β1 enhances legumain activity, secretion, and activation of internalized extracellular prolegumain, and in return, legumain mediates the inhibitory effect of TGF-β1 on osteoblast maturation. Our findings not only advance our understanding of the complex regulatory networks governing osteoblast function but also provide novel insight into the mode of action of TGF-β1 in regulation of skeletal homeostasis. Thus, our studies may unveil potential therapeutic avenues for bone-related disorders mediated by aberrant TGF-β1 signaling.

## 3. Material and methods

## 3.1. Cell culturing

Human bone marrow derived mesenchymal stem cells (hBMSC) overexpressing the human telomerase reverse transcriptase (hBMSC-TERT (Simonsen et al., 2002); hereby referred to as BMSC) were used and cultured in basal medium containing Minimal Essential Media (MEM) with L-glutamine, 10% (v/v) fetal bovine serum (FBS), 1% penicillin (100 U/mL) and streptomycin (100 µg/mL). Human embryonic kidney 293 (HEK293; American Type Culture Collection (ATCC), Rockville, USA) and monoclonal legumain over-expressing HEK293 (M38L) cells were cultured as previously described (Smith et al., 2012). HEK293 and M38L cells were seeded at a density of 5 × 10^4^ cells/cm^2^, maintained at 37°C and 5% CO^2^ in a humidified atmosphere before serum-free conditioned media (CM) were collected after 4 days and used as reagents for treatment of BMSC.

For osteogenic differentiation, BMSC cells were seeded at a density of 2 x 10^4^ cells/cm^2^ and at 80% confluence, the cells were incubated for 3, 7, or 14 days in 2X osteogenic induction medium (OBIM) (Jafari et al., 2017) diluted 1:1 with conditioned media (CM) from M38L (prolegumain concentration 165 ng/mL; measured by ELISA) or HEK293 (control, prolegumain concentration 1.1 ng/mL; ELISA) cells. In addition, the BMSC cells were incubated with or without 25 ng/mL TGF-β1 (R&D systems, MN, USA) and/or 50 µM of the irreversible legumain inhibitor RR-11a analog (referred as RR-11a; MedChemExpress, NJ, USA). An equal volume of solvent (4 nM HCl and 3% bovine serum albumin in distilled H_2_O (dH_2_O) for TGF-β1 and DMSO for RR-11a) was used as control. The medium was changed every 3-4 days.

### 3.2. Harvesting of conditioned media and cell lysates

Cell conditioned media were collected, centrifuged at 800 rpm for 10 minutes at 4°C and frozen at -20°C. Adherent cells were washed with PBS before adding legumain lysis buffer (100 mM sodium citrate, 1 mM disodium-EDTA, 1% n-octyl-β-D-glucopyranoside, pH 5.8) or Buffer RLT Pluss buffer (QIAGEN, Hilden, Germany). Cell lysates were sonicated for 20 seconds and centrifuged at 10 000 g for 5 minutes before the supernatants were frozen at -20°C (in legumain lysis buffer), -70°C (in Buffer RLT) or directly analyzed. Total protein concentration in cell lysates was measured at 595 nm according to (Bradford, 1976) and the manufacturer (Bio-Rad Laboratories, Hercules, CA) in a microplate reader (Wallac Victor® Nivo™, Perkin Elmer, Boston, MA, USA). Bovine serum albumin (0-400 µg/mL) was used as a standard for calculation of total protein concentrations. All measurements were performed in triplicates.

### 3.3. Immunoblotting and enzyme-linked immunosorbent assay (ELISA)

Gel electrophoresis and immunoblotting were performed by applying 15 µg total protein using NuPAGE 4-12% gels (Life Technologies) and the supplied NuPAGE MOPS SDS running buffer, prior to transfer to a nitrocellulose membrane (Trans-Blot® Turbo™ Mini-size nitrocellulose) in the Trans-Blot® Turbo™ Transfer System for 30 minutes. The membranes were blocked for 1 hour at room temperature with Odyssey® Blocking Buffer and probed with goat polyclonal anti-legumain (1:200, AF2199, R&D systems) or mouse monoclonal anti-GAPDH (1:10 000, MAB5718, R&D systems) antibody in blocking buffer/0.2% T-TBS (Tween 20 in TBS; 1:1) overnight at 4°C. Membranes were subsequently washed 3-4 times in T-TBS buffer and incubated with donkey anti-goat IR Dye 680LT (1:10 000, LI-COR, Cambridge, UK) or donkey anti-mouse 800CW (1:10 000, LI-COR) for 1 hour at room temperature. After another washing procedure, membranes were briefly dried and analyzed using Odyssey-CLx Imaging System (LI-COR). Immunoband intensities were quantified using the Image Studio Lite 5.2 software (LI-COR).

Legumain concentration in conditioned media was measured using a total legumain ELISA kit (DY4769; R&D Systems) according to the manufacturer’s protocol.

### 3.4. Legumain activity assay

Cleavage of the peptide substrate Z-Ala-Ala-Asn-AMC (Bachem, Bubendorf, Switzerland) was used to measure the proteolytic activity of legumain in cell lysates as previously described (Johansen et al., 1999). In brief, 20 µL sample, 100 µL assay buffer (39.5 mM citric acid, 121 mM Na_2_HPO_4_, 1 mM Na_2_EDTA, 0.1% CHAPS, pH 5.8 and 1 mM DTT), 5 µL of the pan-cathepsin protease inhibitor E64 (to abolish unknown asparagine endopeptidase activity, 1 µM final concentration, Sigma Aldrich, MA, USA) and 50 µL peptide substrate solution (final concentration 10 µM) were added in black 96-well microtiter plates (Corning Life Science, MA, USA). Kinetic measurements based on increase in fluorescence (360EX/460EM) over 10 or 60 minutes were performed at 30°C in Victor® Nivo™ microplate reader. Kinetics were calculated as peak increase in fluorescence per second (dF/sec) and corrected for total protein concentration (µg/mL) in the sample.

### 3.5. Quantitative PCR

Total RNA was extracted and purified from cell lysates harvested in Buffer RLT Pluss using a RNeasy® Plus Kit (QIAGEN, Hilden, Germany) according to the manufacturers protocol. Complement DNA (cDNA) was synthesized using High-Capacity cDNA Reverse Transcription Kit (Applied Biosystems, Warrington, UK) and TaqMan Reverse Transcription Reagents (Applied Biosystems, Warrington, UK) in a PerkinElmer 2720 Thermal Cycler (PerkinElmer, Shelton, CT, USA) (25°C for 10 min, 37°C for 90 min, 85°C for 5 min). RT-qPCR was performed using PowerTrack SYBR Green Master Mix (Applied Biosystems, Warrington, UK) and the Applied Biosystems StepOnePlus™ Instrument with the accompanying software StepOne™. β-actin forward (5’-3’: ACC GAG CGC GGC TAC A) and reverse primer sequence (5’-3’: TCC TTA ATG TCA CGC ACG ATT T). GAPDH forward (5’-3’: GTC TCC TCT GAC TTC AAC AGC G) and reverse primer sequence (5’-3’: ACC ACC CTG TTG CTG TAG CCA A). Legumain forward (5’-3’: GCA GGT TCA AAT GGC TGG TAT) and reverse primer sequence (5’-3’: GGA GTG GGA TTG TCT TCA GAG T). ΔCT was calculated as the CT value of the gene of interest, minus the mean of CT values of housekeeping genes (β-actin and GAPDH). ΔΔCT was calculated as ΔCT, minus the average ΔCT of control samples.

### 3.6. Quantification of matrix mineralization

Matrix mineralization was quantified using alizarin red staining as previously described (Jafari et al., 2015). In brief, the conditioned medium was removed before the cells were washed briefly in PBS and fixed in 70% ice-cold ethanol at -20°C for one hour. The fixed cells were rinsed in dH_2_O and stained for 10 minutes with tilting in 40 mM aqueous alizarin red solution (pH 4.2) at room temperature. Excess dye was removed with dH_2_O, followed by three washes in PBS to reduce nonspecific staining. The stained cells were scanned using a photo scanner (Epson Perfection V600, Epson, Suwa, Japan) before alizarin red was eluted in an aqueous 20% methanol and 10% acetic acid solution. Aliquotes (100 µL) of the eluate was added in triplicates to a 96-well plate and absorbance was measured at 570 nm in a microplate reader (Wallac Victor® Nivo™, Perkin Elmer).

Matrix mineralization was also quantified using IRDye® 800CW BoneTag™ (LI-COR, Cambridge, UK) as previously described (Bosnjak et al., 2019). In brief, BoneTag™ (final concentration 2 pmol/mL) was added to the cell culture medium the day before analysis. After 24 hours, the conditioned medium was removed, and the cells were washed with PBS. Fresh PBS was added, and fluorescence was measured at 800 nm using the Odyssey-CLx Imaging System (LI-COR) and quantified using the Image Studio Lite 5.2 software.

### 3.7. Statistics

Data is presented as mean ± standard error of mean (SEM). Two-tailed students T-test, one-way, two-way or three-way ANOVA was performed when appropriate. Statistical significance was considered at p<0.05. Calculations were performed using GraphPad Prism (V9.0; GraphPad Software, Inc., San Diego, CA, USA) or R Statistical Software (V4.2.2; (R Core Team, 2021)).

## 4. Results

### 4.1. TGF-β1 inhibits formation of mineralized matrix by osteogenic BMSC cultures

To investigate the mechanism involved in TGF-β1-induced inhibition of osteoblast maturation, we employed osteogenic cultures derived from a human bone marrow mesenchymal stem (stromal) cell line, referred to as BMSC. To test if this cell model is suitable for our studies, we first determined the effect of TGF-β1 on osteoblast maturation and mineralization, by culturing osteogenic cultures of BMSC with or without TGF-β1 (25 ng/mL) for 14 days. Formation of mineralized matrix in the osteogenic cultures was assessed by two methods, using quantification of BoneTag™ fluorescence (Fig. 1A) and alizarin red staining (Fig. 1B) and showed TGF-β1-induced inhibition of mineralization and osteoblast maturation.

**Figure 1.**
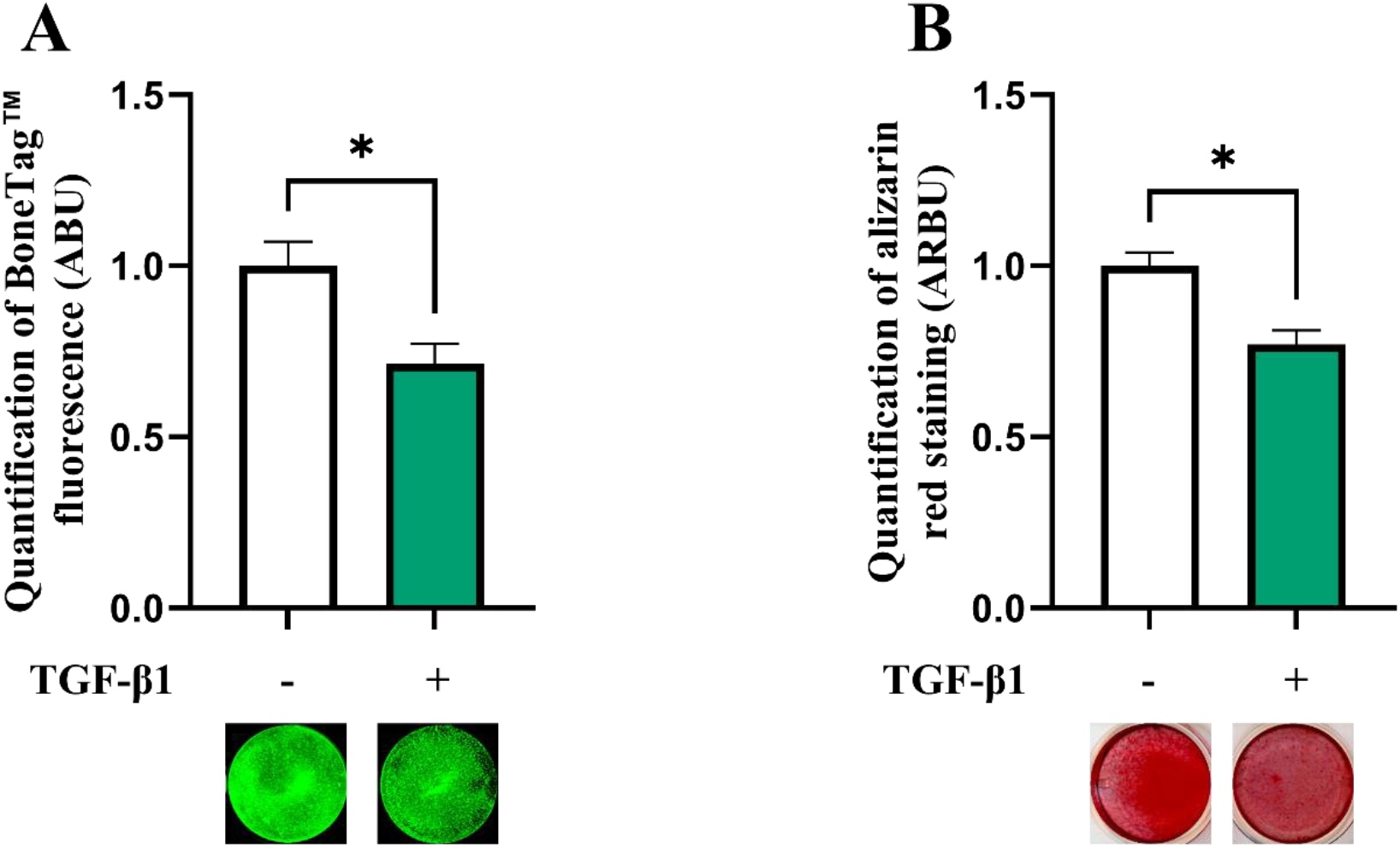
TGF-β1 inhibits mineralized matrix formation in osteogenic BMSC cultures. BMSC were cultured in osteogenic induction medium with or without TGF-β1 (25 ng/mL) for 14 days. Matrix mineralization was measured by BoneTag™ fluorescence (**A**, n=4) and alizarin red staining (**B**, n=3). Normalized data represent mean ± SEM. Two-tailed unpaired Student’s t-test. *p<0.05. Numbers (n) represent individual biological replicates.

### 4.2. TGF-β1 enhances legumain activity and secretion, but not mRNA expression in osteogenic BMSC cultures

We have reported legumain as a novel inhibitor of osteoblast maturation (Jafari et al., 2017). To assess whether the inhibitory effect of TGF-β1 could be mediated through altered legumain expression and function in osteogenic cultures, legumain was analyzed in BMSC cultures in the presence or absence of TGF-β1 (25 ng/mL) during osteogenic differentiation for up to 14 days. Immunoblot analysis of cell lysates from osteogenic cultures on day 3, 7, and 14 of differentiation showed that TGF-β1 did not significantly affect protein expression of either pro-or mature legumain at any time point, although a tendency towards increased levels of prolegumain was observed on day 14 of differentiation (Fig. 2A-C). We then determined the effect of TGF-β1 on the activity of endogenous legumain in osteogenic cultures and observed increased legumain activity in cells cultured with TGF-β1 for 14 days (Fig. 2D). In addition, treatment with TGF-β1 increased the concentration of legumain in the conditioned media at day 3 and 7, indicating increased legumain secretion (Fig. 2E). In contrast, legumain mRNA expression was decreased by TGF-β1, reaching significance at 7 days of osteogenic differentiation (Fig. 2F).

**Figure 2.**
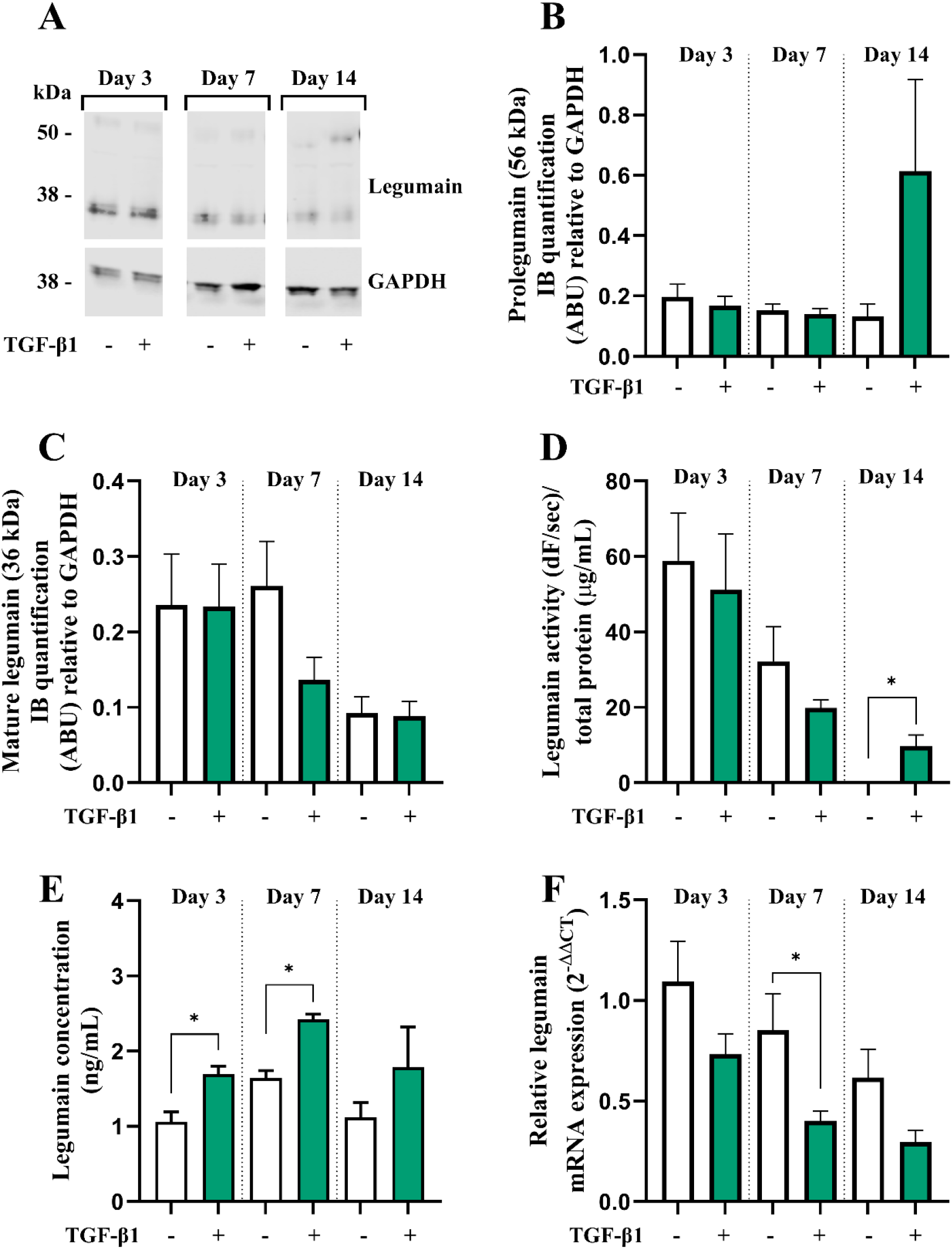
TGF-β1 enhances legumain activity and secretion, but not mRNA expression in osteogenic BMSC cultures. BMSC were cultured in osteogenic induction medium containing 1:1 conditioned control medium (HEK293-CM) with or without TGF-β1 (25 ng/mL) for 3, 7 or 14 days. **A**. One representative immunoblot of legumain and GAPDH (housekeeping) in cell lysates (n=4). **B-C**. Quantification of the 56 kDa (**B**; prolegumain) and 36 kDa (**C**; mature legumain) immunoband (IB) intensities as arbitrary units (ARBU) relative to GAPDH on immunoblots presented in A (n=4). **D**. Legumain activity (dF/sec) in cell lysates adjusted for total protein concentration (µg/mL) (n=3). **E**. Legumain concentration (ng/mL) in conditioned media measured by ELISA (n=6). **F**. Legumain mRNA expression relative to the mean CT values of two housekeeping controls (GAPDH and β-actin) (2^-ΔΔCT^; n=4-6). **B-F:** Data represent mean ± SEM. **B-E:** Two-tailed students t-test. **F**. Two-tailed students t-test on ΔCT values. *p<0.05. Numbers (n) represent individual biological replicates.

### 4.3. TGF-β1 enhances activation of internalized prolegumain during osteogenic differentiation of BMSC

Prolegumain has previously been shown to be secreted by various cell types, to be present in extracellular environments (e.g. plasma) and to be internalized and activated by recipient cells (Lunde et al., 2019, Smith et al., 2012). Since we demonstrated that legumain also was secreted during osteogenic differentiation and that the secretion was enhanced by TGF-β1, we aimed to investigate whether prolegumain could be internalized and activated by BMSC. The cells were incubated in a 1:1 mixture of basal medium and conditioned media from HEK293 cells overexpressing and secreting legumain (prolegumain concentration 165 ng/mL; (Smith et al., 2012)) or HEK293 control cells (control; prolegumain concentration 1.1 ng/mL) for 24 hours before the media was switched to fresh basal medium. Cell lysates were harvested 1, 2, 3, 4 and 6 days after prolegumain exposure. Immunoblot analysis of cell lysates showed that prolegumain (56 kDa) was internalized and almost completely processed to mature legumain (36 kDa) 24 hours after exposure to extracellular prolegumain, whereas the legumain level was back to baseline after 4 days. (Fig. 3A). We also found significantly increased legumain activity in BMSC lysates 1 and 2 days post exposure (DPE) to prolegumain containing media, whereas legumain activity levels returned to baseline 3 DPE. (Fig. 3B).

**Figure 3.**
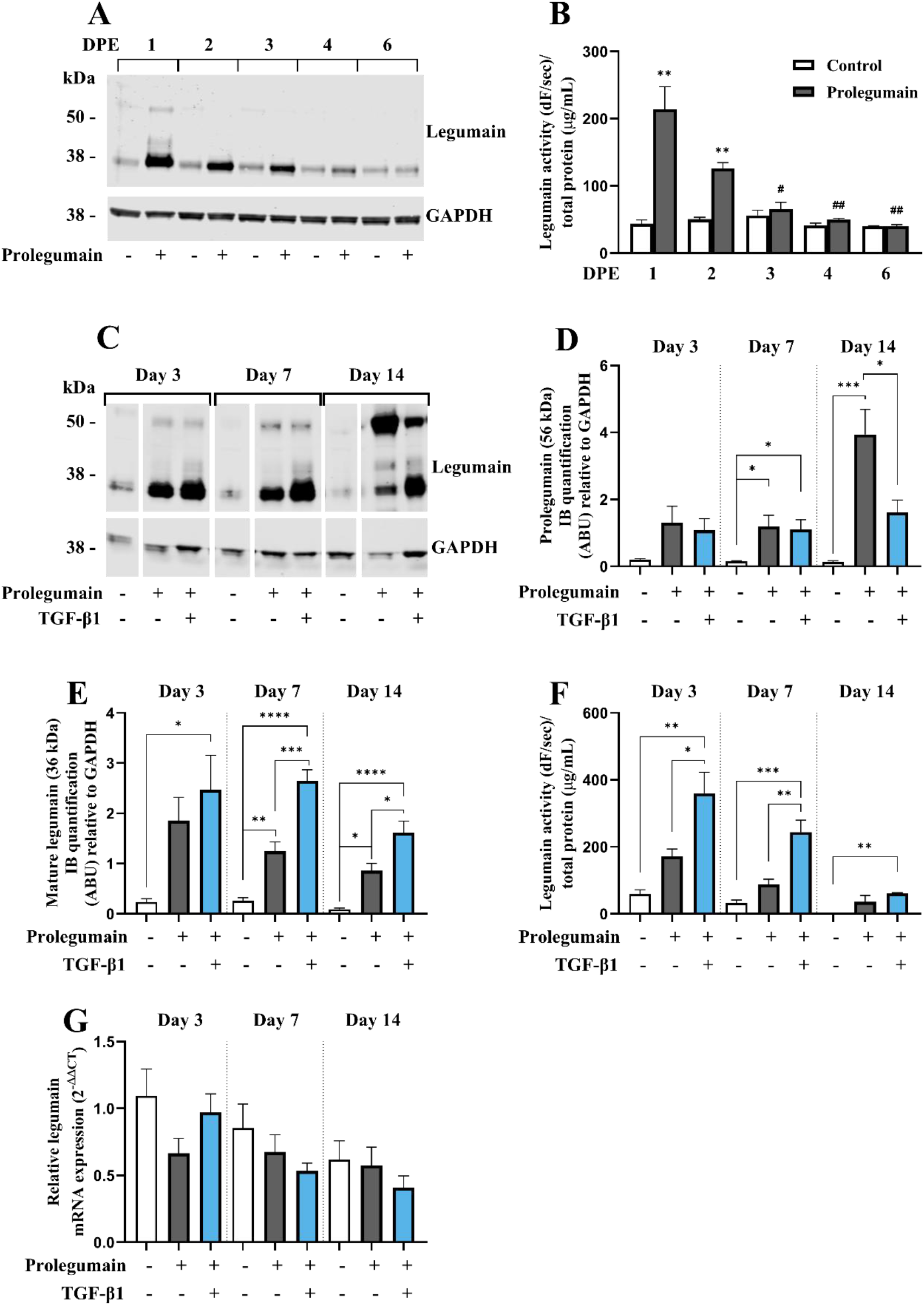
TGF-β1 enhances activation of internalized prolegumain during osteogenic differentiation. **A-B**. BMSC were cultured in basal medium containing 1:1 HEK293-conditioned medium with (165 ng/mL) or without (control) prolegumain for 24 hours before the medium was changed to basal medium. Lysates were harvested 1, 2, 3, 4 and 6 days post exposure (DPE). **C-G**. BMSC were grown in osteogenic induction medium containing 1:1 HEK293-conditioned medium with (165 ng/mL) or without prolegumain and with or without TGF-β1 (25 ng/mL) for 3, 7 or 14 days. **A, C**. One representative immunoblot of legumain and GAPDH (housekeeping) in cell lysates (n=3-4). **D-E**. Quantification of the 56 kDa (**D**; prolegumain) and 36 kDa (**E**; mature legumain) immunoband (IB) intensities as arbitrary units (ARBU) relative to GAPDH on immunoblots presented in C (n=4). **B, F**. Legumain activity (dF/sec) in cell lysates adjusted for total protein concentration (µg/mL) (n=3). **G**. Legumain mRNA expression relative to the mean of CT values of two housekeeping controls (GAPDH and β-actin) (2^-ΔΔCT^; n=4-6). **B, D-G:** Data represent mean ± SEM. **B:** Two-tailed students T-test. **D-F:** Two-way ANOVA (Šidák correction). **G**. Two-way ANOVA (Šidák correction) on ΔCT values. *p<0.05, **p<0.01, ***p<0.001, ****p<0.0001 (**B**: compared to control on same DPE). #p<0.05, ##p<0.01 compared to same treatment on 1 DPE. Numbers (n) represent individual biological replicates.

Next, we investigated whether prolegumain could be internalized by osteogenic cells at different stages of differentiation, and if TGF-β1 regulated legumain internalization and activation. BMSC were cultured in osteogenic induction medium with or without prolegumain-rich medium and with or without TGF-β1 (25 ng/mL) for 3, 7 or 14 days. Immunoblot analysis of cell lysates revealed significantly increased level of prolegumain (56 kDa) at day 7 and 14, whereas mature legumain (36 kDa) was increased at all time points upon exposure to prolegumain, supporting internalization and processing of prolegumain in osteogenic cultures (Fig. 3C-E). Interestingly, concomitant treatment with prolegumain and TGF-β1 significantly reduced the cellular level of prolegumain at day 14. In contrast, increased level of mature legumain was observed at day 7 and 14 by combined treatment with TGF-β1 and prolegumain, compared to prolegumain treatment alone. In addition, we observed a tendency towards increased legumain activity in osteogenic cultures exposed to extracellular prolegumain, whereas combined treatment with TGF-β1 and prolegumain significantly increased legumain activity on day 3 and 7 of osteoblast differentiation. (Fig. 3F). We did not observe any changes in legumain mRNA expression upon exposure of osteogenic cultures to extracellular legumain and/or TGF-β1 (Fig. 3G). In summary, these results indicated that TGF-β1 promotes processing and activation of internalized legumain.

### 4.4. Legumain can inhibit osteoblast maturation through a paracrine mechanism that is exacerbated by TGF-β1

After establishing that secreted prolegumain can be internalized and activated during osteogenic differentiation, we examined if internalized prolegumain could act as a paracrine factor to inhibit matrix mineralization and if such inhibition was regulated by TGF-β1. BMSC were cultured for 14 days in osteogenic differentiation medium with or without legumain-rich condition media and with or without TGF-β1 (25 ng/mL). BoneTag™ fluorescence (Fig. 4A) and alizarin red staining (Fig. 4B) showed decreased matrix mineralization by treatment with prolegumain alone. In addition, concomitant treatment with prolegumain and TGF-β1 showed additive inhibition of mineralization.

**Figure 4.**
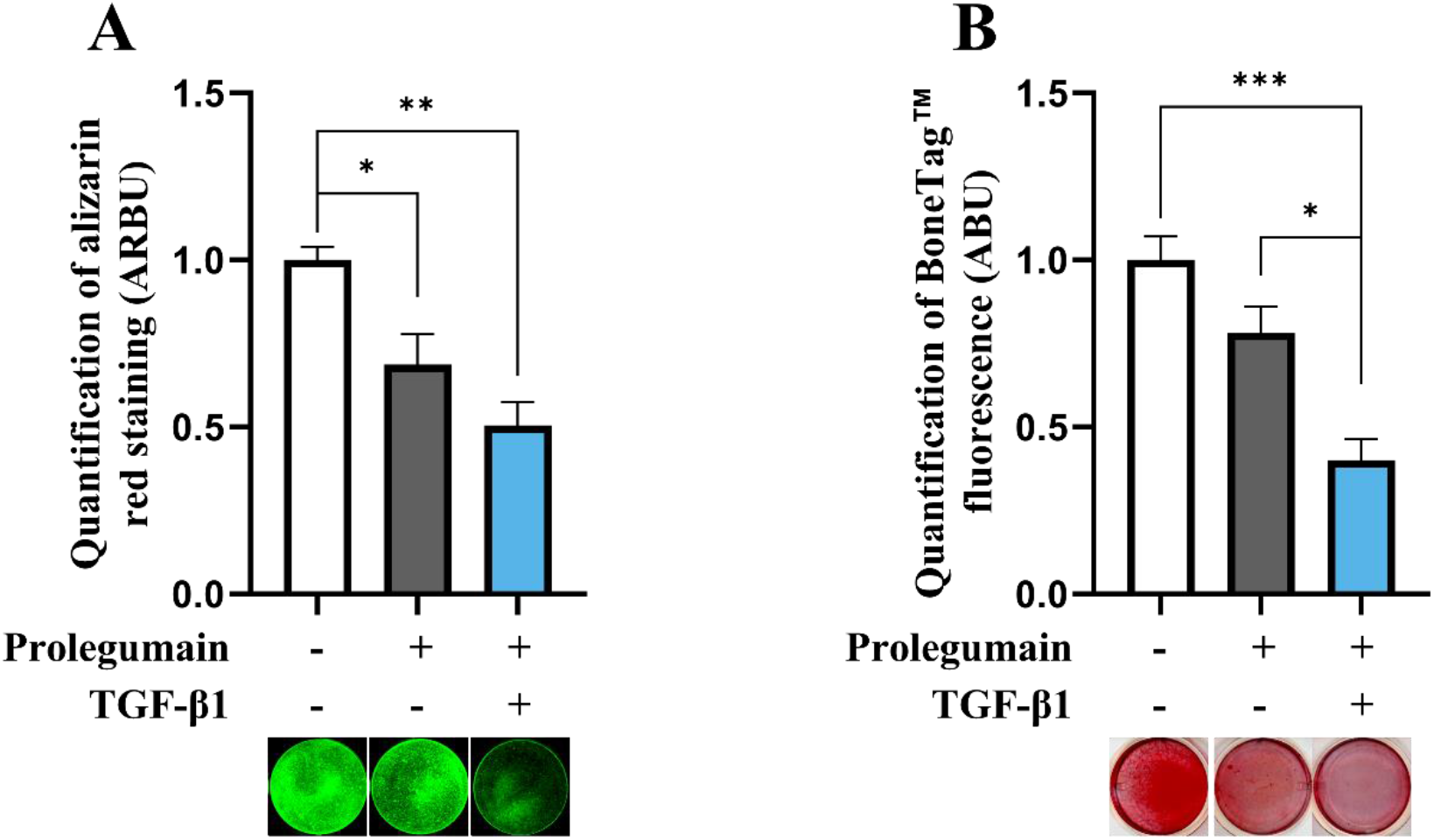
Treatment with legumain inhibits matrix mineralization and is exacerbated by concomitant treatment with TGF-β1. BMSC were cultured in osteogenic induction medium with (165 ng/mL) or without prolegumain-rich conditioned medium alone or with TGF-β1 (25 ng/mL) for 14 days. Normalized matrix mineralization measured by BoneTag™ fluorescence (**A**, n=4) and alizarin red staining (**B**, n=3). Data represent mean ± SEM. Two-way ANOVA (Šidák correction). *p<0.05, **p<0.01, ***p<0.001. Numbers (n) represent individual biological replicates.

### 4.5. Reduced osteogenic mineralization by prolegumain treatment is abolished by legumain inhibition

Although presence of TGF-β1 in osteogenic BMSC cultures increased legumain levels, associated with inhibited osteoblast maturation, this association does not necessarily prove that legumain mediates the inhibitory effect of TGF-β1 on osteoblast maturation. Therefore, we employed a selective, irreversible legumain inhibitor (RR-11a) and showed that presence of RR-11a (50 µM) in osteogenic BMSC cultures on day 14 significantly inhibited legumain activity, in the presence of legumain-rich media alone or together with TGF-β1 (Fig. 5A). As expected, RR-11a abolished the inhibitory effect of extracellular legumain on osteoblast maturation (Fig. 5B). Interestingly, the TGF-β1-mediated inhibition of osteoblast maturation was completely abolished upon pharmacological inhibition of legumain activity by RR-11a. This data provides direct experimental evidence for the role of legumain in mediating the inhibitory effect of TGF-β1 on osteoblast maturation.

**Figure 5.**
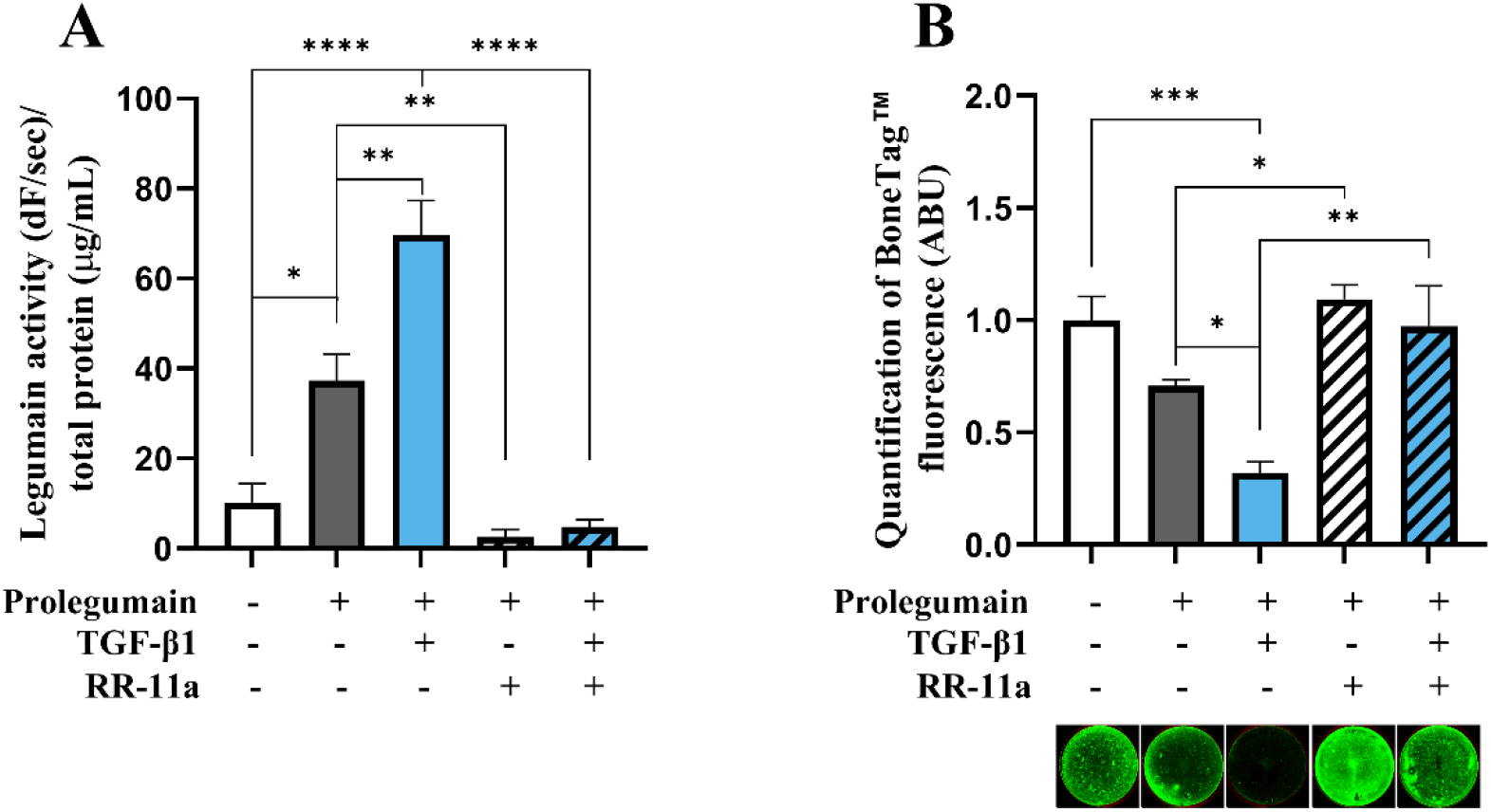
Pharmacological inhibition of legumain rescue matrix mineralization in osteogenic cells treated with TGF-β1 and prolegumain. BMSC were differentiated to osteogenic cells for 14 days with or without treatment with 1:1 prolegumain-rich conditioned medium, TGF-β1 (25 ng/mL) and/or the legumain inhibitor RR-11a (50 µM). **A**. Legumain activity (dF/sec) in cell lysates adjusted for the total protein concentration (µg/mL) (n=3-4). **B**. Normalized matrix mineralization measured by BoneTag™ fluorescence. Data represent mean ± SEM (n=3-4). Three-way ANOVA (Tukey’s post-hoc). *p<0.05, **p<0.01, ***p<0.001, ****p<0.0001.

## Discussion

Our study reveals a novel regulatory mechanism of osteogenesis, indicating that TGF-β1 modulates osteoblast maturation through legumain. Highly expressed in bone, TGF-β1 is believed to be synthesized and deposited in the bone matrix in a latent form by osteoblasts during bone formation. It is then released and activated during bone resorption, playing a pivotal role in coupling bone resorption with formation. Extensive literature on the effects of TGF-β1 on skeletal homeostasis presents diverse and occasionally conflicting conclusions, as reviewed by (Janssens et al., 2005) and (Wu et al., 2016). This variability in findings can be partially attributed to differences in experimental design and conditions across various research groups. Despite these differences, it is broadly accepted that TGF-β1 exerts a pleiotropic influence on osteogenic differentiation by promoting proliferation and early commitment to the osteoblastic lineage via non-canonical (Smad-independent) signaling pathways. On the other hand, it has been suggested that TGF-β1 inhibits osteoblast maturation through canonical (Smad-dependent) TGF-β signaling, as detailed by (Wu et al., 2016). While the mechanisms behind the promoting effect of TGF-β1 on early osteoblast differentiation are well-established, the mechanisms underlying its inhibitory effect on late-stage osteoblast differentiation (i.e. osteoblast maturation) remain poorly understood. Our study demonstrates that TGF-β1 regulates legumain at different levels to modulate osteoblast maturation. We observed that TGF-β1 enhanced legumain activity and secretion in osteogenic cultures, despite a decrease in legumain mRNA expression. This suggests a possible post-transcriptional regulation of legumain by TGF-β1, potentially involving protein stabilization or increased processing of endogenous legumain to the mature form. We observed that osteogenic cells can internalize and activate extracellular prolegumain, leading to increased intracellular legumain activity. Interestingly, TGF-β1 treatment promoted the processing and activation of internalized prolegumain.

We have previously shown that legumain inhibits osteoblast maturation through autocrine mechanism (Jafari et al., 2017). The discovery that legumain is present in the secretome of various cell types and exists within extracellular compartments has spurred speculation about its potential paracrine regulatory roles (Lunde et al., 2019). Here, we show that extracellular legumain is internalized and activated at the early stages of osteogenic differentiation, similar to what we have previously described in kidney (HEK293) cells (Smith et al., 2012), although the legumain protein levels and activity return to baseline in a time-dependent manner. We observed that extracellular prolegumain inhibited osteoblast maturation, evidenced by decreased formation of mineralized matrix. Therefore, our findings provide novel experimental evidence that legumain acts as a paracrine regulator of osteoblast differentiation and function.

Interestingly, we observed that extracellular prolegumain and TGF-β1 exert an additive effect on inhibition of osteoblast maturation. This observation was in line with our findings that TGF-β1 increases legumain secretion, internalization, and activation, but did not necessarily prove that legumain mediates the inhibitory effect of TGF-β1 on osteoblast maturation. Therefore, we used pharmacological inhibition of legumain activity in osteogenic cultures and observed that the inhibitory effect of extracellular prolegumain and TGF-β1 was completely abolished upon legumain inhibition. This interesting observation provides strong evidence for the inhibition of osteoblast maturation by legumain being mediated through proteolytic activity and for the role of legumain in mediating the inhibitory effect of TGF-β1 on osteoblast maturation.

In the present study, we observed that TGF-β1 promotes the activation of internalized legumain during osteoblast differentiation. In most contexts, the process and significance of the internalization and subsequent activation of lysosomal proteases remain largely unexplored. TGF-β1 has been reported to promote autophagy in several cancer types (He et al., 2019, Kiyono et al., 2009), presumably in a Smad and JNK-dependent manner (Suzuki et al., 2010). In addition, increased lysosomal activity has been demonstrated to have an important role in the cellular remodeling during epithelial-to-mesenchymal transition (EMT) (Kern et al., 2015), an oncogenic process of which TGF-β1 is a major driver. Therefore, it is possible that the promoting effects of TGF-β1 on legumain secretion, internalization, and activity, contribute to promotion of autophagy and increased lysosomal activity observed in EMT.

Our study provides novel insight into the complex regulatory network governing osteoblast maturation unveiling a novel regulatory axis involving TGF-β1 and legumain. By demonstrating the role of legumain in TGF-β1-mediated inhibition of osteoblast maturation, our study has important implications for bone disorders associated with aberrant TGF-β1 signaling and suggests legumain as a potential target for therapeutic interventions. Future studies are warranted to further explore the detailed mechanisms by which TGF-β1 regulates legumain secretion and function. Additionally, investigating the *in vivo* relevance of this pathway using animal models of bone diseases will be crucial for assessing its translational potential.

## 6. Conclusion

Legumain was shown as a paracrine regulator of osteoblast maturation. In addition, increased activation of endogenous and exogenous legumain was demonstrated by TGF-β1. An additive inhibitory effect on osteoblast maturation was shown by concomitant treatment with TGF-β1 and exogenous prolegumain. Complete rescue of mineralization phenotype was demonstrated by pharmacological inhibition of legumain in cells concomitantly treated with prolegumain and TGF-β1, providing evidence that the inhibitory effect of TGF-β1 is mediated through legumain activation.

## 8. Acknowledgement

Hilde Nilsen is highly acknowledged for the technical assistance. Rigmor Solberg is a member of the COST action CA20113 ProteoCure (A sound proteome for a sound body: targeting proteolysis for proteome remodeling).

